# A report on a characteristic vocalization in *Corvus macrorhynchos osai* with an indication of vocal learning

**DOI:** 10.1101/2021.09.21.461160

**Authors:** Noriko Kondo

**Affiliations:** Rikkyo University

**Keywords:** vocal learning, corvid, crow, subsong, babbling, dialect, vocal development

## Abstract

In the process of vocal learning, animals give immature vocalization, such as babbling and subsong. Here I report a characteristic vocalization given by a subspecies of large-billed crow, *Corvus macrorhynchos osai*, that live in Kuroshima Island, Okinawa, Japan. This vocalization (type-K) is characterized by very rapid bill movements (ca. 13 times/sec). The type-K was heard throughout the island, indicating that the population of the Kuroshima Island share this call type. In addition, a juvenile crow was observed giving immature type-K repeatedly. This observation suggests that this call type is not innate but acquired through vocal learning.

## Introduction

Vocal production learning, or vocal learning, is the ability to produce non-innate vocalization through social learning and supposed to be essential for human language (Jarvis 2006). In the animal kingdom, this ability can be found only in limited taxa: songbird, hummingbird, and parrot in birds, as well as cetaceans, elephants, bats, and pinnipeds in mammals (Petkov and Jarvis 2012).

In vocal learning animals, young individuals acquire new vocalization by practicing it repeatedly in the course of development. This practicing behavior is called “babbling” in humans (Lenneberg 1967). The babbling behavior is also reported in other vocal learning mammals, such as pygmy marmosets *Cebuella pygmaea* (Elowson et al. 1998) and sac-winged bats *Saccopteryx bilineata* (Knörnschild et al. 2006). In songbirds, the immature song produced by the juveniles during the sensorimotor phase is called “subsong”. Subsong is soft, highly variable, and rambling vocalization, lacking species-specific characteristics in the early stages. As the birds develop, the subsong would become more stable, but still acoustically different from the adult song. This stage of song is referred to as “plastic song”. During the subsong and plastic song period, the juveniles are supposed to learn the motor skill to produce sounds that would match their vocal template, which is learned during the sensitive phase (Nottebohm 1970, Nottebohm 1972, Wilbrecht and Nottebohm 2003).

Vocal learning often results in the variation of vocalizations between geographically distant populations. The geographic variation of vocalization is also referred to as vocal cultures or “dialects” (Catchpole and Slater 2008). The dialect in animals was first reported in songs of white-crowned sparrow *Zonotrichia leucophrys* (Marler and Tamura 1964) and is now widely reported in songbirds. Recent studies have revealed the geographic variation also exists in bird and mammal calls, although calls have been believed to be innate and lack flexibility compared to songs (Boughman and Moss 2003).

As corvids belong to Passeriformes, it is not surprising that they have vocal learning abilities. Their vocal learning ability is reported in some studies as well as lots of anecdotes, although they do not produce songs comparable to those of songbirds. Webber and Stefani (1990) reports A captive California scrub jay *Aphelocoma coerulescens superciliosa* learned calls of Florida scrub jays *A. c. coerulescens*. In ravens *Corvus corax* it is reported that their vocal repertoire is supposed to be formed via cultural transmission, especially within sexes (Enggist-Dueblin and Pfister, 2002). New Caledonian crows, *Corvus moneduloides*, give calls acoustically different depending on the distance (Bluff et al. 2010). Corvids under captivity are also reported to learn human vocalizations, suggesting their flexible vocal learning ability (Chamberlain and Cornwell, 1971; Bluff et al. 2010).

Large-billed crow *Corvus macrorhynchos* is known anecdotally to learn human vocalizations, indicating their vocal learning ability, but no scientific publication exists (see Karasawa 1988 for anectdotes, for example). In addition, whether they acquire their species-specific vocal repertoire through vocal learning is not clear. In a study of the Indian population, the acoustic structure is different among subspecies but how such acoustic differences emerge is not known (Martens et al. 2000). Fledglings of large-billed crows give various unstable vocalizations and Kuroda (1969) considered this vocalization was functionally comparable to the subsong of songbirds. However, no study on vocal development in large-billed crows exists so far, especially focusing on a particular call type.

Here, I report a preliminary observation of vocal learning in a wild population of *Corvus macrorhynchos osai*, a subspecies of large-billed crow. *C. m. osai* live in Yaeyama Islands, Okinawa. They are smaller in size than *C. m. japonensis*, a subspecies of large-billed crow that lives in the mainland of Japan. The groups that live in different islands are morphologically different, especially in bill length, depending on the habitat they live in (Yamasaki 2010). However, their vocal behavior is not reported at all thus far. Here I report vocal behavior in *C. m. osai*, especially focusing on a specific vocalization that is not reported in any published literature of vocal behavior in large-billed crows.

## Study site

The Study site was Kuroshima Island (24° 14’ 22” N, 124°0’53” E), one of Yaeyama Islands of Okinawa Prefecture, Japan. The island is very flat (>10m above sea level) since it was formed when the coral reef rose. It is very small in size and has an area of about 10.02 km^2^, with a peripheral length of 12.62 km. Since the main industry of this island is cattle-breeding, the island is for the most part covered with grassland. Although the original habitat of large-billed crows is forest (Goodwin 1986), *C. m. osai* in Kuroshima Island seems to have highly adapted to the open grassland (Yamasaki 2010).

## Methods

From September 4^th^ to 7^th^ 2013 and from January 23^rd^ to 27th 2014, I visited Kuroshima Island and recorded vocalization of *C. m. osai*. I searched for vocalizing crows by bicycle, and as soon as I spotted vocalizing crows, I started recordings using a directional microphone (MKH416-P48U3, Sennheiser) and a linear PCM recorder (DR-100MKII, Tascam). The recordings were made from 0700-1830. From Oct. 22^nd^ through Oct. 26th of 2020, I again visited Kuroshima Island and plotted the places where the vocalization was heard onto a map. Unfortunately, however, I could not record their vocalizations using a microphone due to the bad weather condition (rain and strong wind). I, instead, tried to capture their vocalization via a digital camera (OM-D E-M1 Mark II, Olympus) using its video recording function.

## Results

In 2013, I noticed very strange vocalization given by the crows of Kuroshima (Figure 1, Video S1). This vocalization consists of two phases: rattling phase and rapid-bill-movement phase. The rattling phase is the introductory phase consisted of a mechanical rattling sound. For the five vocalizations I managed to record, the rattling phase lasted for 220-577 msec. In *C. m. japonensis* this phase is referred to as “rattling call”, which is given independently (Kuroda 1990). The following rapid-bill-movement phase was 849-1048 msec. In this phase, the crows move their bills very rapidly (ca. 13 times/sec) while vocalizing, resulting in a vibrato sound. When giving this call, the crow’s head directed downwards (see Video S1). This call type was also recorded in 2014. This call type is not described in any publications, so I named it “type-K call” (K after Kuroshima Island). Unfortunately, however, it was very difficult to record this call since the crows did not give this vocalization very frequently. Thus, I only managed to record one call in 2013 and four calls in 2014.

**Figure1.**
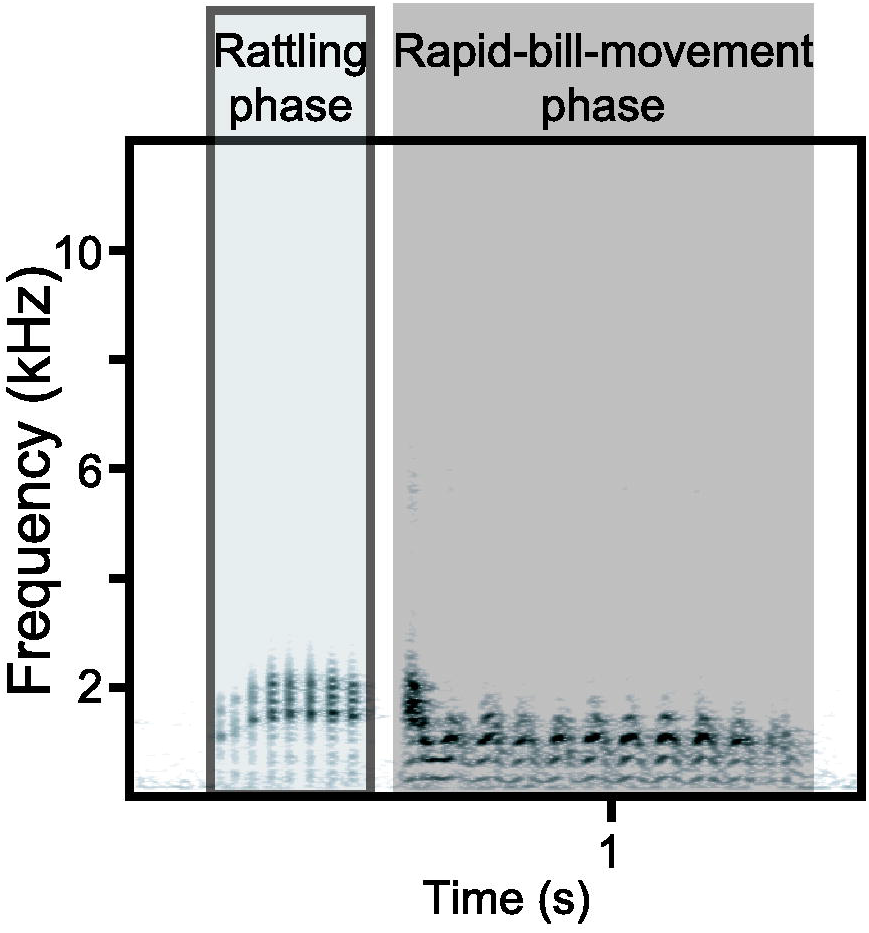
Sound spectrogram of type-K call given by an adult bird. The call is composed of a rattling phase (gray area with dark gray outline) and a rapid-bill-movement phase (highlighted in dark gray).

In 2020, the type-K was heard in 15 places of the island (Figure 2). At one of these points, I observed one crow giving immature vocalization. This bird was young and considered to be born that year, judging from its pink mouth cavity (Kitagawa 1980). It was wandering around cattle in front of a cattle shed while vocalizing. Its vocalization contained an introductory rattling phase as well as adults’ type-K, but the following rapid-bill-movement phase was replaced by several short notes of various duration and frequencies (Figure 3, Video S1). The rattling phase was also produced independently, resulting in a rattling call, but the short notes were always preceded by the rattling phase. In Kuroshima Island I had never heard any vocalization composed of rattling phase and other sounds, except for type-K. Therefore, I judged this juvenile’s vocalization as immature type-K. This bird repeatedly gave similar vocalization, that is, rattling call followed by various short notes, for more than five minutes after I found it. It then flew into the cattle shed and I could not follow it anymore.

**Figure2.**
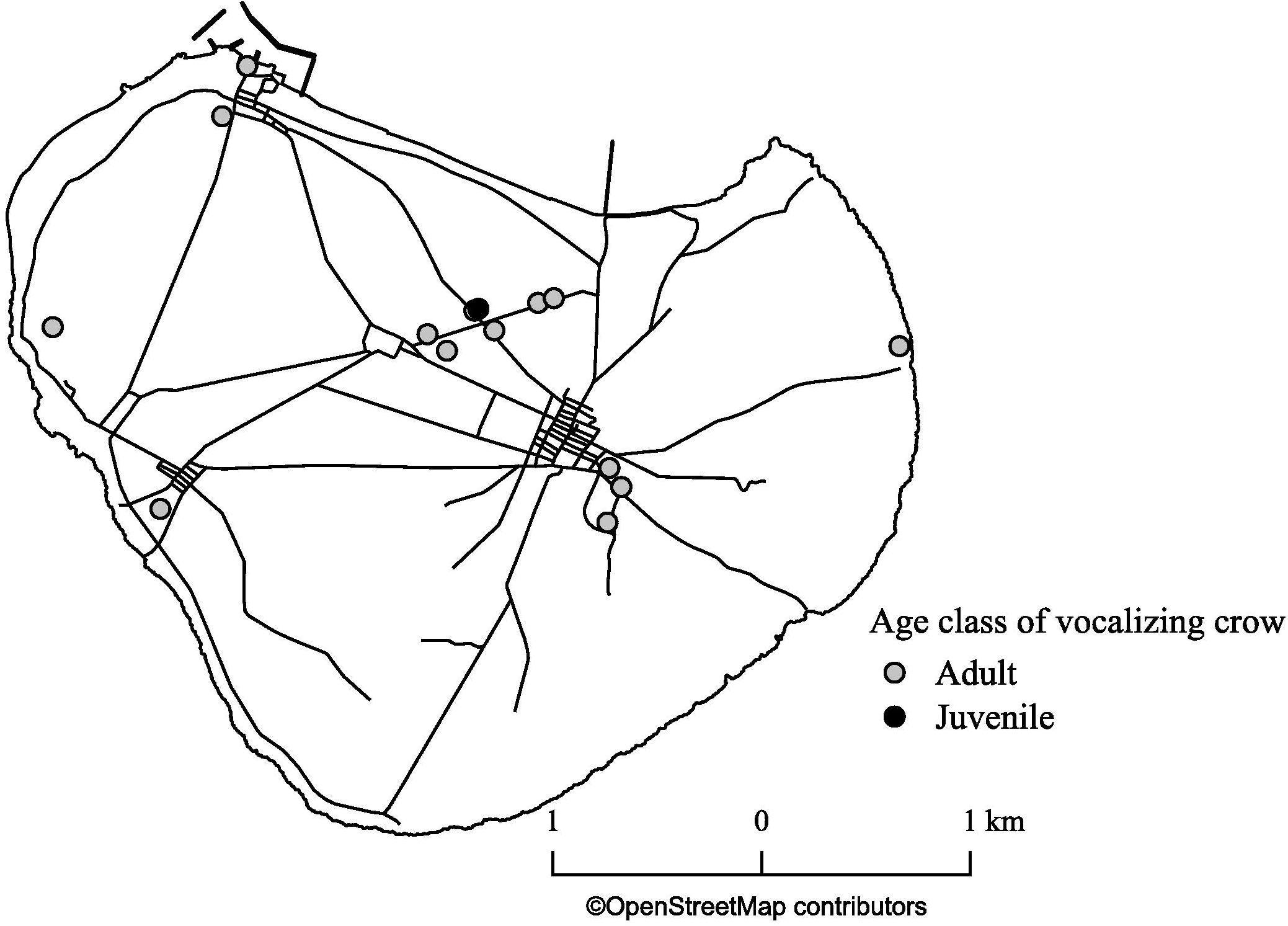
A map of Kuroshima Island showing the points where type-K was recorded. The grey circles indicate the points where adult type-K calls were recorded and the black shows the point where a juvenile crow gave immature type-K.

**Figure3.**
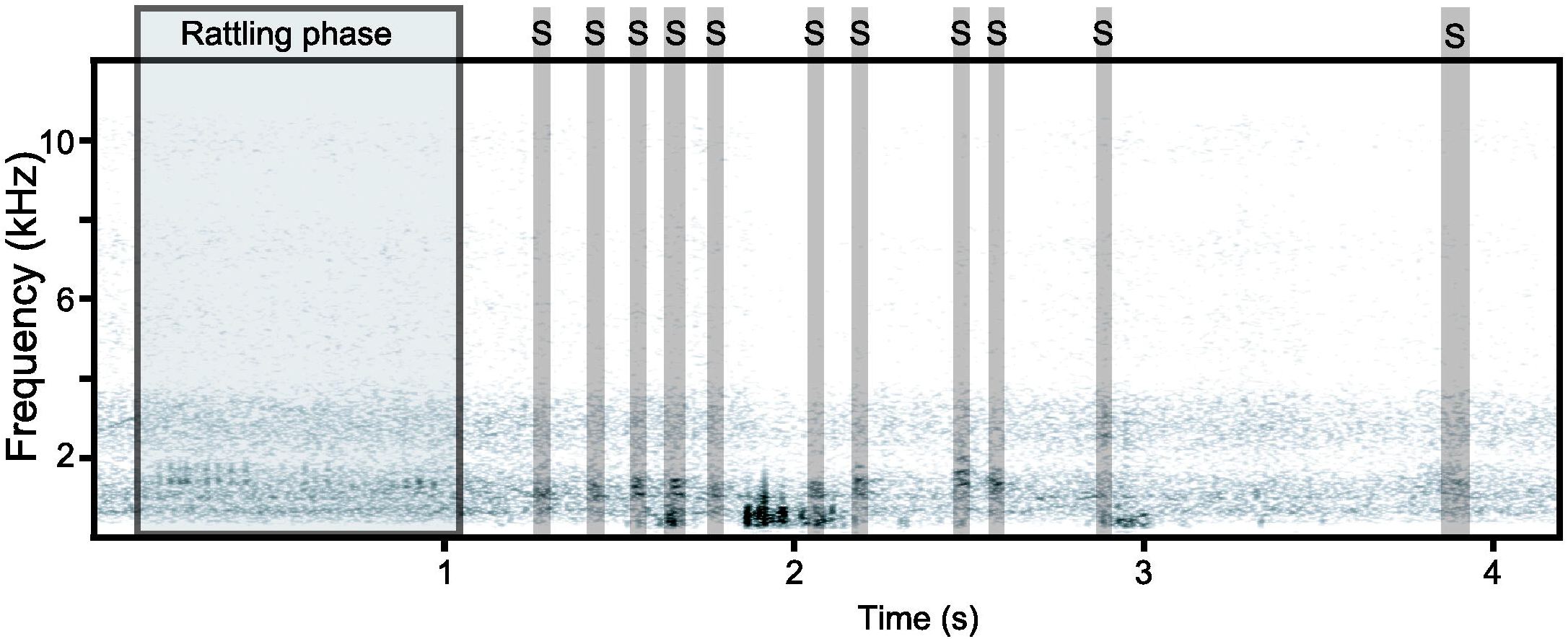
Sound spectrogram of immature type-K call given by a juvenile bird. The juvenile immature call was composed of rattling phase (gray area with dark gray outline) and repetitive short notes indicated by the dark gray highlights and the capital Ss. Note that the rapid-bill-movement phase in adult call was replaced by short notes in juvenile vocalization, and the intervals, as well as lengths of short notes, were not stable. The sound spectrogram was created after high-pass filtering (0.3 kHz).

## Discussion

The unique vocalization, type-K, has never been reported in any literature of vocalization of *C. m. japonensis*, suggesting that this call is at least subspecies-specific. In addition, the juvenile immature vocalization implies that this call is not innate but learned. It is not surprising that the *C. m. osai* show vocal learning ability since it is known anecdotally that the large-billed crow learns anthropologic sound and human speech. Still, this is the first report showing the juvenile large-billed crows produce subsong-like immature vocalizations of a particular call type, indicating that their vocal repertoire is at least to some extent acquired through vocal learning.

In the rapid-bill-movement phase of adult type-K, it is supposed that they produce one single sound while moving bills very rapidly, resulting in the vibrato-like frequency modulation, since the sounds of the rapid-bill-movement phase are not separated but connected. In the juvenile type-K, on the other hand, the rapid-bill-movement was replaced by a sequence of short notes. It may be physically demanding for the juveniles to move the bill rapidly (at the speed of ca. 13 times/sec) while vocalizing. The juvenile might have imitated the rapid-bill-movement phase by repeatedly giving short notes since the adult rapid-bill-movement phase sounds like a very rapid repetition of short notes. As I only observed one case in this study, it is not clear how juvenile crow acquires rapid-bill-movement as it develops. How type-K changes as the crow develops would reveal the acquisition of complex body movement related to their vocalization.

The function of type-K is not clear at all because most of the crows giving type-K in my observation were just sitting on a branch alone without showing any particular behavior. From the acoustic profile, I speculate that the type-K may function in the same way as the rattling call since it contains a rattling sound in the introductory phase. The function of rattling call is also unknown, but it is relatively soft vocalization and thus supposed to be used in short distance communication. Kuroda (1990) reported that he had heard the rattling call during fights. In one case of my observation, one crow was repeatedly giving type-K while snatching feed mixture from the cattle inside a cowshed, together with more than 15 crows. Although I could not observe them closely due to the distance, social interaction including fights over the food may have occurred between them. If this was the case, it would support the idea that the type-K and rattling call are similar in their function. The head downward vocalizing posture of type-K also indicates that this call is not for long-distance communication, since such posture would disturb sound propagation. How they use this call in their social interaction would be revealed in future studies of long-term and close observation of *C. m. osai*.

Type-K call seemed to be shared by several individuals within the island. As I have not identified the individual identities of the vocalizing crows, the vocalizations by the same individuals may have been recorded repeatedly. In addition, given the small island size, it is not difficult for crows to fly between the farthest points where type-K had been heard. However, it is not natural to suppose that a single type-K calling crow moved around all over the island as I moved. Rather, it is reasonable to suppose that several individuals of this island give type-K. In fact, I had heard two individuals gave type-K call at the same time. The observation of a practicing juvenile also supports the idea that the type-K is shared within the island population through vocal learning.

I do not, however, know whether this call is also shared with populations outside of Kuroshima Island. I preliminarily recorded vocalization of *C. m. osai* also in Ishigakijima Island, Iriomotejima Island, Kohamajima Island, and Haterumajima Island. In these four islands, type-K was not recorded. This may be due to the fact the degree of difficulty to record crows’ vocalization differed according to the islands because of the size differences. Kuroshima Island is the smallest among them, and therefore easy to record various vocalizations of *C. m. osai*. In order to find out whether this call type is shared with other islands or not, a lot of thorough recording on other islands is strongly required.

As the bird song dialect is supposed to be related to the site fidelity (Lemon 1975), the dispersion from the islands should also be considered. However, it is not clear whether the crows of the Yaeyama Islands disperse to other islands or not. The distance between the islands is within ca. 20 km (the distance from Kuroshima Island to Ishigakijima Island is ca. 16 km, Kuroshima Island to Iriomotejima Island is ca. 10 km, with Aragusuku Island in between). Given that the daily flight distance of *C. m. japonensis* of the urban area is ca. 7km in the non-breeding season (Morishita et al. 2004), it is not impossible for *C. m. osai* to fly to other islands. Still, Yamasaki (2010) points out the possibility of speciation of *C. m. osai* within the Yaeyama Islands based on their behavioral and morphological differences. This suggests that the crow population of each island does not disperse frequently. As the dialect in bird song is considered to play important role in speciation (Slabbekoorn and Smith 2002), the study of vocalization might uncover the speciation within the *C. m. osai*.

The study of vocal behavior of large-billed crows is unfortunately not flourishing. This study is only descriptive, but I believe it is suggesting. I hope much more studies would be conducted in the future to understand vocal communication in large-billed crow, a highly social species.

## Supporting information

Electric Supplementary Video

## Acknowledgment

The research trips to Yaeyama Islands in 2013 and 2014 were financially supported by JSPS KAKENHI Grand Number JP13J05437. I am grateful to my human family for always let me follow crows freely.

## Electric Supplementary Materials

Video S1. A video of an adult and a juvenile crow giving type-K.

